# Temperature Stressed *Caenorhabditis elegans* Males Prioritize Feeding over Mating Resulting in Sterility

**DOI:** 10.1101/2022.03.01.482587

**Authors:** Nicholas B. Sepulveda, Lisa N. Petrella

**Affiliations:** Marquette University

## Abstract

Heat stress negatively impacts fertility in sexually reproducing organisms at sublethal temperatures. These temperature stress effects are typically more pronounced in males. In some species, sperm production, quality, and motility are the primary cause of male infertility under temperature stress. However, this is not the case in the model nematode *Caenorhabditis elegans* where changes in mating behavior are the primary cause of fertility loss. We report that temperature stressed *C. elegans* males experience a dramatic upset in the balance of their food drive and their mating drive such that they prioritize feeding over mating. This change in priorities is due partially to increased expression of the chemoreceptor *odr-10* in the AWA sensory neurons. Increased *odr-10* expression in the presence of ample food demonstrates that males are unable to experience satiety, thus they never leave a food source and engage in mate searching behavior. These results demonstrate that moderate temperature stress may have profound and previously underappreciated effects on reproductive behaviors. As climate change associated temperature variability becomes more commonplace, it will be imperative to understand how temperature stress affects conserved behavioral elements critical to reproduction.

## Introduction

Sexual reproduction and its consequent genetic recombination offer a multitude of advantages for organisms that employ it (Crow, 1994; Otto and Lenormand, 2002; Barton, 2009). However, it has an ancient and conserved weakness: Temperature sensitivity. Species as diverse as Asian rice, fruit flies, nematodes, cattle, and humans all exhibit a dramatic loss of fertility at temperatures just above their optimal fertility temperature range (Lam and Miron, 1996; David et al., 2005; Prasad et al., 2006; Petrella, 2014; Morrell, 2020). While female fertility can be negatively impacted by temperature stress, male fertility is often more vulnerable and contributes disproportionately to infertility in sexually reproducing organisms (David et al., 2005; Zinn et al., 2010; Petrella, 2014; Iossa, 2019; Walsh et al., 2019a). In ectothermic organisms that cannot regulate their body temperature, like the nematode *Caenorhabditis elegans*, temperature effects on fertility are even more pronounced (Kingsolver et al., 2013; Walsh et al., 2019b). These organisms frequently play important roles as nutrient recyclers in their ecosystems, and *C. elegans* is no exception (Queirós et al., 2019). As climate change-associated temperature uncertainty increases, the necessity of understanding precisely how these changes affect the fertility of important organisms like *C. elegans* is crucial.

Prior research has often focused on the negative impact of temperature stress on germline-associated processes in males, such as sperm production and quality in metazoans like fruit flies, cattle, and humans (Casady et al., 1953; Hjollund et al., 2002; David et al., 2005; Ahmad et al., 2012; Rao et al., 2016; Green et al., 2019; Llamas-Luceño et al., 2020). However, we have previously shown that in temperature stressed *C. elegans* males the impact of temperature stress on germline-associated processes like sperm production and activation is relatively minor (Nett et al., 2019), and cannot account for the near total sterility observed (Petrella, 2014; Sepulveda and Petrella, 2021). Rather, we found that in *C. elegans*, male sterility under temperature stress is largely defined by negative impacts on somatic-associated processes, specifically mating organ anatomy and mating behavior (Nett et al., 2019). Similar phenomena have been observed in *Drosophila* males, which experience reduced reproductive ability under temperature stress that can be attributed to negative changes in behavior (Fasolo and Krebs, 2004; Dolgin et al., 2006; Jørgensen et al., 2006). As behavioral defects that arise under temperature stress limit reproduction even when germline function is not unduly impacted, understanding the negative effects of temperature on behavior merits priority.

*C. elegans* serves as an ideal model system to dissect the negative impact of temperature stress on male reproductive behavior. *C. elegans* male mating behaviors are sophisticated and many components of these behaviors are understood on the cellular and genetic level (Barrios, 2014; Chute and Srinivasan, 2014; Barr et al., 2018; Emmons, 2018; Muirhead and Srinivasan, 2020). Male *C. elegans* behavior is generally dictated by two necessities: mating and feeding. Balancing these priorities is crucial to successful reproduction. The ideal environment for *C. elegans* males in the wild or the lab is one where food and mates are plentiful and spatially overlapping. However, if there are no hermaphrodites, a male will engage in mate searching behavior and leave food to locate a mate. Males sense potential mates by both hermaphrodite excreted chemical signals and touch (Chute and Srinivasan, 2014; Barr et al., 2018; Muirhead and Srinivasan, 2020). The best understood chemical signaling molecules in *C. elegans* are the ascarosides (Butcher, 2019; Park et al., 2019; Muirhead and Srinivasan, 2020). Males perceive hermaphrodite ascaroside messages via the male-specific sensory neurons (CEM neurons) in the head as well as chemosensory neurons shared with hermaphrodites (AWA, AWC, ADF, and ASK), and will accumulate in the vicinity of these molecules (Srinivasan et al., 2008; Izrayelit et al., 2012; Narayan et al., 2016; Fagan et al., 2018). Sensing a hermaphrodite by touch is the more potent stimulus, which males accomplish using male-specific RnA/RnB neurons in the rays of their tail (Barrios, 2014; Barr et al., 2018). An adult male on a food source that has not experienced physical contact with a hermaphrodite will dramatically alter their locomotion resulting in food leaving in search of a mate (Barrios, 2014). Our previous work has shown both that male tail anatomy and the ability of males to respond the presence of hermaphrodites are negatively impacted under temperature stress (Nett et al., 2019), thus sensory and other male reproductive behaviors could be negatively impacted in a similar fashion.

Here, we asked which components of *C. elegans* male mating behavior are negatively impacted by temperature stress. We find that temperature stress (27°C) profoundly changes decision making in *C. elegans* males by upsetting feeding/mating prioritization: Temperature stressed males prioritize food over mating. The cumulative result of this change in priorities is that very few temperature stressed males are interested in mating and thus very few reproduce successfully. Our findings show that temperature stress can elicit underappreciated and dramatic changes in behavior with profoundly negative consequences for reproduction.

## Methods

### Strains used

*Caenorhabditis elegans* worms were maintained using standard methods at 20°C on nematode growth media (NGM) plates (Brenner, 1974) spotted with 200 μl AMA1004 *Escherichia coli*. Males of wild isolate strains were generated by heat shocking L4 hermaphrodites at 30°C for five hours followed by recovery at 20°C and selection of male progeny. Male populations were subsequently maintained by crossing males and hermaphrodites of the same genotype at 20°C. JU1171, LKC34, N2, and MT7929 *[unc-13(e51)]* were obtained from the *Caenorhabditis* Genetics Center (Minneapolis, MN, USA) which is funded by NIH Office of Research Infrastructure Programs (P40 OD010440). UR460 *[him-5(e1490); kyIs53 (Podr-10::odr-10::GFP)]* was obtained from the Portman lab (Rochester NY, USA). The wild isolate strains used in this study were isolated from the following localities: JU1171 (Concepción, Chile), LKC34 (Madagascar), and N2 (Bristol, UK).

### Temperature treatments

Two temperature treatments were used in this study: (1) Continuous exposure to 20°C, and (2) Continuous exposure to 27°C. 20°C experiments were performed on male worms maintained continuously at 20°C. 27°C experiments were performed on male worms upshifted to 27°C at the L1 stage and maintained continuously at 27°C until assayed. For mate search experiments, *unc-13(e51)* hermaphrodites were maintained at 20°C, and only exposed to elevated temperature at the young adult stage for the duration of experiments at 27°C. For both temperature treatments, male celibacy was ensured by isolating males from hermaphrodites at the L4 stage and allowing them to mature to adulthood overnight (18-24 hours). Males were isolated at a population density of ≤ 20 worms per plate.

### Pheromone response assay

Pheromone response assays were performed similarly to Sammut et al. (2015). NGM plates were prepared two days before the experiment. One day before the experiment plates were spotted with 50 μl AMA1004 *E. coli* grown to OD_600_ = 1.0. One day before the experiment, males and strain matched (same genotype) hermaphrodites were isolated from each other at the L4 stage and allowed to mature to adulthood (18-24 hours). On the day of the experiment, adult hermaphrodites were rinsed twice with M9 buffer dispensed in a three-well watch glass to remove residual *E. coli*. These adult hermaphrodites were used to make “hermaphrodite-conditioned buffer” by placing them in the lid of a 1.7 ml microfuge tube with M9 buffer at a concentration of one hermaphrodite per microliter of M9 for five hours in a humidity chamber at 20°C. After the conditioning period, the buffer was removed and stored at 4°C in a new 1.7 ml microfuge tube until use within two hours. Test plates were prepared by dispensing 1 μl plain M9 buffer and 1 μl conditioned M9 buffer across from each other, 2 mm from the center of the bacterial food spot. Experiments were performed after the buffer spots had completely dried (three to five minutes). Five to eight adult test males were rinsed twice with M9 buffer dispensed in a three-well watch glass to remove residual *E. coli*. and placed on the test plate equidistant from the control and the conditioned spots, 3 mm away from the center. Males were recorded at 1X magnification using a Nikon SMZ 1500 stereo microscope equipped with an HR Plan APO 1X objective lens (Nikon Instruments Inc., Melville NY, USA) for 20 minutes using a Q imaging Exi Blue camera (Teledyne Photometrics, Tucson AZ, USA) and Q Capture Pro 7 software (Teledyne Photometrics, Tucson AZ, USA). Three to five test plates per strain and temperature condition were imaged for each experiment, with a single 20-minute video captured per plate. Five biological replicates were performed for each strain and temperature condition, with n ≥ 15 males scored total. Videos were subsequently analyzed using FIJI software (Schindelin et al., 2012). Each time a male entered either the control or conditioned spot, the amount of time the male spent in the spot was scored. Each entrance to a spot was considered an independent event *if* after leaving, the male worm had moved at least 2 mm away from the spot before re-entering. If males encountered each other in either spot and engaged in mating behavior for at least five seconds, the corresponding entrance events were discarded from the analysis. Data were analyzed with a two-way ANOVA using Tukey’s multiple comparison test in Prism 9 (Graphpad, San Diego, CA, USA).

### Leaving assay

Leaving assays were performed in accordance with Lipton et al. (2004). NGM plates were prepared two days before the experiment. One day before the experiment plates were spotted with 18 μl AMA1004 *E. coli* grown to OD_600_ = 1.0. Plates were used the day after spotting without exception. Two days before the experiment, strain matched (same genotype) hermaphrodites were isolated from males at the L4 stage and allowed to mature to adulthood. One day before experiments the adult hermaphrodites were fixed in a fresh 4% formaldehyde in PBS solution overnight at 4°C (Barrios et al., 2008). On the day of experiments formaldehyde-fixed (FA) hermaphrodites were washed three times in PBS buffer. Test plates were prepared by placing two FA hermaphrodites equidistant from each other 2 mm from the center of the food spot. Control plates were prepared by disturbing the food lawn at the location where each hermaphrodite would be placed on an experimental plate. One day before the experiment, males were isolated from hermaphrodites at the L4 stage and allowed to mature to adulthood. The day of the experiment, a single male was rinsed once in M9 buffer and was placed at the center of the food spot of an experimental or control plate. Plates were examined at intervals (2 hours, 4 hours, 6 hours, 8 hours, 20 hours, and 24 hours) to check for “leavers”. A male was considered a leaver and left censored if the male or worm tracks were observed on the agar within 2.5 mm of the edge of the plate. Ten test plates and ten control plates per strain were set up for each experiment and four replicates were performed with a n = 40 total. We used the previously published model of male leaving behavior (Lipton et al., 2004), which employs a single exponential decay function which remains constant over the course of an experiment: N(t)/N (0)=exp(-λt), [where N(t) refers to the number of males retained at a given time and the hazard value (λ) gives an estimate for the probability of leaving (P_Leaving_). P_Leaving_ values were calculated by fitting the data using the R survival package with an exponential parametric survival model using maximum likelihood (Hilbert and Kim, 2017; R Core Team, 2019). The average P_Leaving_ was calculated for each replicate and the data were compared with a two-way ANOVA using Tukey’s multiple comparison test in Prism 9 (Graphpad, San Diego CA, USA).

### Motility assay (crawling on NGM agar plates)

One day before the experiment NGM plates were spotted with 50 μl AMA1004 *E. coli* grown to OD_600_ = 1.0. The bacteria were spread over the NGM plate using a sterile glass spreader to 2 mm from the edge of the plate. The day before the experiment males and hermaphrodites were isolated from each other at the L4 stage and allowed to mature to adulthood. The day of the experiment, ten males or hermaphrodites were washed once in M9 buffer and then placed at the center of the food spot. At least two, but as many as five plates (per sex, strain, and temperature treatment) were set up for each experiment. Subsequently, the males or hermaphrodites were allowed to move about the plate and acclimate for two hours. After the acclimatization period, the plates were imaged at 1X magnification using a Nikon SMZ 1500 stereo microscope equipped with an HR Plan APO 1X objective lens (Nikon Instruments Inc., Melville NY, USA) capturing 30s videos using a Q imaging Exi Blue camera (Teledyne Photometrics, Tucson AZ, USA) and Q Capture Pro 7 software (Teledyne Photometrics, Tucson AZ, USA). Each plate was imaged at least once but as many as six times to capture each worm on the plate. Care was taken to film each worm once, but occasional reimaging of a worm may have occurred. To assess absolute movement velocity and convert pixels to mm, a known mm scale was imaged and recorded. Videos were processed using FIJI; and maximum movement velocities for each worm were calculated using the WrmTrck plugin for FIJI (Schindelin et al., 2012; Nussbaum-Krammer et al., 2015). Detailed instructions for utilizing WrmTrck are available at the WrmTrck website (http://www.phage.dk/plugins/wrmtrck.html). When assessing movement velocity, worms that did not move at all during the 30s time frame or moved less than five seconds (frames) during the 30s period were excluded from the analysis. To assess the proportion of worms that were motile during the 30s test period, worms that did not move at all or moved less than 1 mm from their initial position were considered not moving. When assessing maximum velocity, two to five plates of worms were assessed for each strain, sex, and temperature condition, with four biological replicates being executed for a total n ≥ 50. Maximum velocity was analyzed using a two-way ANOVA using Tukey’s multiple comparison test, and the number of worms in the test population that moved versus those that did not move were analyzed using a Fisher’s exact test in Prism 9 (Graphpad, San Diego CA, USA).

### Physical endurance assay (thrashing in buffer)

Vigor assays were performed in M9 buffer as in Chatterjee at al. (2013). A single male or hermaphrodite was allowed to crawl on the agar surface of an NGM plate to remove excess bacteria and then transferred to M9 buffer in a three well watch glass. The number of full body bends (thrashes) were recorded for 60s. At least three biological replicates were performed for each strain, sex, and temperature condition for n ≥ 60 total. Data were analyzed with a two-way ANOVA using Tukey’s multiple comparison test in Prism 9 (Graphpad, San Diego CA, USA).

### Food search assay

Food search assays were executed similarly to Ryan et al. (2014). One day before the experiment NGM plates were spotted with 7 μl AMA1004 *E. coli* grown to OD_600_= 2.5. One day before the experiment males were isolated from hermaphrodites at the L4 stage and allowed to mature to adulthood. The day of the experiment, a single male was rinsed once in M9 buffer and was placed 3.2 cm away from the food spot. Plates were examined every fifteen minutes for two hours to ascertain if a male had reached the food spot. Ten test plates per strain were set up for each experiment and at least five biological replicates were performed for an n = 50 total. Survival curves were generated using the Kaplan-Meier method and analyzed using the Log-rank (Mantel-Cox) test in Prism 9 (Graphpad, San Diego CA, USA).

### Polystyrene microsphere entrapment assay

Test plates were prepared as for food search assays described above. The day of the experiment, a 20 μl droplet of 1 μm diameter Polybead® Carboxylate Red Dyed Microspheres (Polysciences Inc, Warrington PA, USA; Catalog number 19119-15) in distilled water at a concentration of 1.5167×10^10^ particles/ml (a 1:2 dilution of polystyrene microspheres to water) was placed at the site of male worm release and allowed to dry. Experiments were then conducted as food search assays described above. Survival curves were generated using the Kaplan-Meier method and analyzed using the Log-rank (Mantel-Cox) test in Prism 9 (Graphpad, San Diego CA, USA).

### Mate search assay

Mate search assays were performed as published by Fagan et al. (2018). One day before the experiment, NGM plates were spotted with 10 μl AMA1004 *E. coli* grown to OD_600_ = 1.0 which was spread into a 1 cm x 2.5 cm rectangular food lawn using a sterile glass spreader. One day before the experiment males were isolated from strain matched (same genotype) hermaphrodites at the L4 stage and allowed to mature to adulthood. For experiments using formaldehyde-fixed (FA) hermaphrodites: Hermaphrodites were isolated two days before experiments, and one day before experiments the adult hermaphrodites were fixed in a fresh 4% formaldehyde in PBS solution overnight at 4°C (Barrios et al., 2008). On the day of the experiment, three FA hermaphrodites were rinsed three times in PBS buffer and placed equidistant from each other at one end of the food lawn. Experiments with live *unc-13(e51)* hermaphrodites: One day before the experiment, L4 *unc-13(e51)* hermaphrodites were isolated and allowed to mature to adulthood. On the day of the experiment three *unc-13(e51)* hermaphrodites were rinsed once in M9 buffer, placed equidistant from each other at one end of the food lawn and allowed to condition the plate for two hours. For both experiments: A single male was rinsed once in M9 buffer and placed on the opposite end of the food lawn. Plates were checked every five minutes for one hour to determine if males had reached the vicinity of the hermaphrodites and commenced mating behaviors. Ten plates per strain and temperature condition were set up. At least six biological replicates were performed, with an n ≥ 60 total. Survival curves were generated using the Kaplan-Meier method and analyzed using the Log-rank (Mantel-Cox) test in Prism 9 (Graphpad, San Diego CA, USA).

### Mate/Food prioritization assays

One day before the experiment test plates and control plates were spotted with two 18 μl AMA1004 *E. coli* grown to OD_600_ = 1.0 spots 1.5 cm apart from each other flanking the center of an NGM plate. Two days before the experiments strain-matched (same genotype) L4 hermaphrodites were isolated and allowed to mature to adulthood. One day before experiments the adult hermaphrodites were fixed in a fresh 4% formaldehyde in PBS solution overnight at 4°C (Barrios et al., 2008). The day of the experiment, formaldehyde-fixed (FA) hermaphrodites were rinsed three times in PBS buffer. Test plates were prepared by placing either two FA hermaphrodites equidistant from each other near the center of one food spot, 2 mm from the center. Control plates were prepared by disturbing the food lawn at the location where each hermaphrodite would be placed on an experimental plate. One day before the experiment, males were isolated from hermaphrodites at the L4 stage and allowed to mature to adulthood. The day of the experiment, a single male was rinsed once in M9 buffer and was placed 1.5 cm from the center of an experimental plate or control plate. Ten test plates and ten control plates per strain were set up for each experiment. Plates were examined at intervals (2 hours, 4 hours, 6 hours, 8 hours, 20 hours, and 24 hours) to check for the location of the male. For experimental plates locations were scored as: food spot without hermaphrodites, food spot with hermaphrodites, or agar. For control plates, locations were scored as: left food spot, right food spot, or agar. Each time a male changed location on the plate (e.g., from a food spot without hermaphrodites to a food spot with hermaphrodites), was scored as a “switch”. Data were compared with a twoway ANOVA using Tukey’s multiple comparison test in Prism (Graphpad, San Diego CA, USA).

### Fluorescent microscopy

Visualization and quantification of ODR-10::GFP was performed as in Wexler et al (2020). Two days before imaging, UR460 males and hermaphrodites were isolated from each other at the L4 stage and allowed to mature to adulthood (18-24 hours). To image well-fed animals, young adults were left on standard NGM plates for an additional 24 hours before imaging. To image starved animals, young adults were rinsed 3X in M9 buffer and transferred to fresh unspotted NGM plates with 200 μg/ml ampicillin (to inhibit bacterial growth) for 24 hours before imaging. Worms were mounted on 5% agarose pads and immobilized with 1 mM levamisole in M9 buffer (Manjarrez and Mailler, 2020) and imaged using a Nikon Eclipse TE2000-S inverted microscope equipped with a Plan Apo 60×/1.25 numerical aperture oil objective (Nikon Instruments Inc., Melville NY, USA). Images were captured using a Q imaging Exi Blue camera (Teledyne Photometrics, Tucson AZ, USA) and Q Capture Pro 7 software (Teledyne Photometrics, Tucson AZ, USA). Scoring of ODR-10::GFP fluorescent intensity was done using a scale of 0-3 where: 0 = absent, 1 = faint, 2 = moderate, and 3 = bright (Ryan et al., 2014; Lawson et al., 2019; Wexler et al., 2020). The scorer was blind to the sex, nutritional status, and temperature treatment of the animal in each image. Data were analyzed using a Chi-square test in Prism 9 (Graphpad, San Diego CA, USA).

## Results

### Temperature stressed males are have a stronger response to hermaphrodite excreted chemical signaling molecules than unstressed males

Because temperature stressed males demonstrate a lack of interest in mating with hermaphrodites (Nett et al., 2019), we hypothesized that these males were unable to detect hermaphrodite chemical signals. To test this possibility, we performed pheromone response assays (Fig. 1A) (Sammut et al., 2015). We predicted that temperature stressed males would spend equal amounts of time in the control region and hermaphrodite conditioned region in comparison to their unstressed counterparts (Simon and Sternberg, 2002). In our experiments we used three wild type strains; N2 the canonical wild type lab strain and two recently isolated wild type strains, JU1171 and LCK34. Unexpectedly, we found that temperature stressed males showed a stronger response to hermaphrodite conditioned buffer than unstressed males (Fig. 1B-D). Temperature stressed males spent more time in the region spotted with hermaphrodite-conditioned buffer than the control region spotted with buffer alone for all three genotypes (Fig. 1B-D). While unstressed LCK34 and N2 males did spend more time in the conditioned region than the control region the time spent was less than the temperature stressed males (Fig. 1 C-D). Surprisingly, unstressed JU1171 males did not spend more time in the conditioned region than the control region (Fig. 1B). These results suggest that *C. elegans* male pheromone response may increase, especially in duration, under temperature stress; however, these males are still mostly sterile and produce very few viable progeny at 27°C (Petrella, 2014; Sepulveda and Petrella, 2021). Notably, *C. elegans* male attraction to hermaphrodite-sourced chemical mating cues at 20°C may not be a universal feature of the species as only LKC34 and N2 males spend more time in the conditioned region at this temperature.

**Fig. 1.**
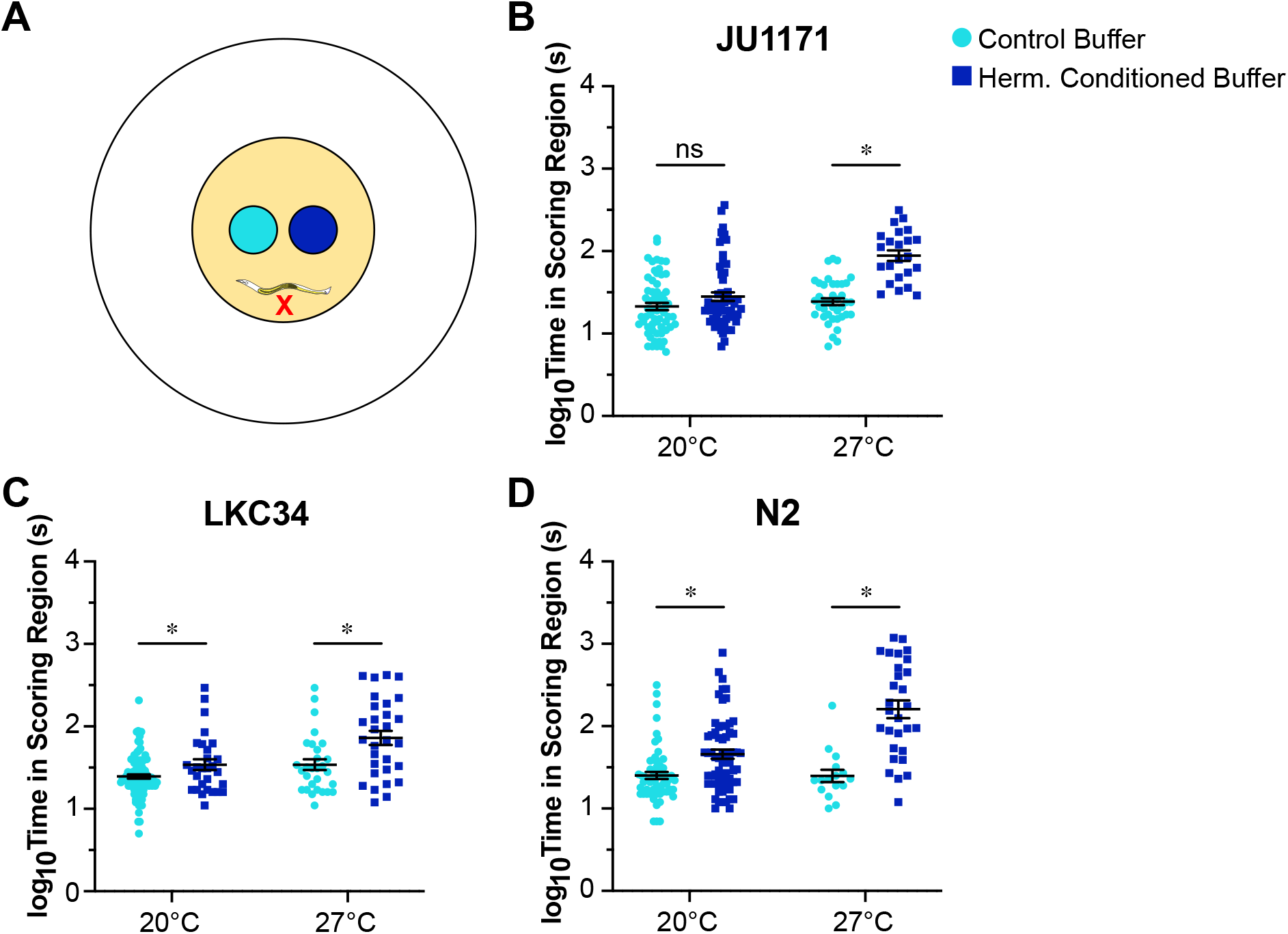
*C. elegans* males exposed to temperature stress (27°C) spend more time in a region conditioned by hermaphrodites than their unstressed (20°C) counterparts. (A) Schematic of the pheromone response assay. NGM plates are spotted with AMA1004 *E. coli* in LB (yellow). The food is spotted with M9 buffer conditioned by strain matched hermaphrodites (dark blue) and M9 alone (light blue). Males were released (red x) and observed for 20 minutes, and their time spent in either region was scored. (B-D) Temperature stressed males of all three strains (JU1171, LKC34, and N2) spend more time in the conditioned region at 27°C. At 20°C, only LKC34 and N2 males spend more time in the conditioned region (Two-way ANOVA with Tukey’s correction, *P ≤ 0.05). Error bars represent ± 1 SEM.

### Temperature stressed males rarely leave a food source even if hermaphrodites are absent

*C. elegans* males use a touch response to detect and respond to hermaphrodites through sensory neurons in the tail fan (Barrios, 2014; Barr et al., 2018). In the absence of hermaphrodites, males at 20°C will leave the food source and engage in mate searching behavior (Lipton et al., 2004; Barrios, 2014). As temperature stressed males have abnormal tail anatomy (Nett et al., 2019), we predicted that temperature stressed males may have an impaired touch response. To test male ability to detect hermaphrodites by touch alone, we fixed hermaphrodites in formaldehyde (FA hermaphrodites) to preserve them and eliminate the production of chemical signaling molecules; and performed leaving assays (Fig. 2A) (Lipton et al., 2004). As expected, unstressed males of all three strains quickly leave a food source without FA hermaphrodites (-FA) but are retained on a food source where they are present (+FA) (Fig. 2B-D). However, we found that temperature stressed males very rarely leave a food source whether FA hermaphrodites are present or not (Fig. 2B-D). To directly compare male leaving behavior under different conditions, we calculated probabilities of leaving (P_Leaving_) from the leaving data (Lipton et al., 2004; Hilbert and Kim, 2017). Temperature stressed males demonstrate very low P_Leaving_ values in the presence or absence of FA hermaphrodites (Fig. 2E-G). Conversely, unstressed males exhibit high P_Leaving_ values in the absence of FA hermaphrodites and low P_Leaving_ values in their presence (Fig. 2E-G). Together, these results suggest that temperature stressed males do not exhibit the characteristic mate searching of their unstressed counterparts.

**Fig. 2.**
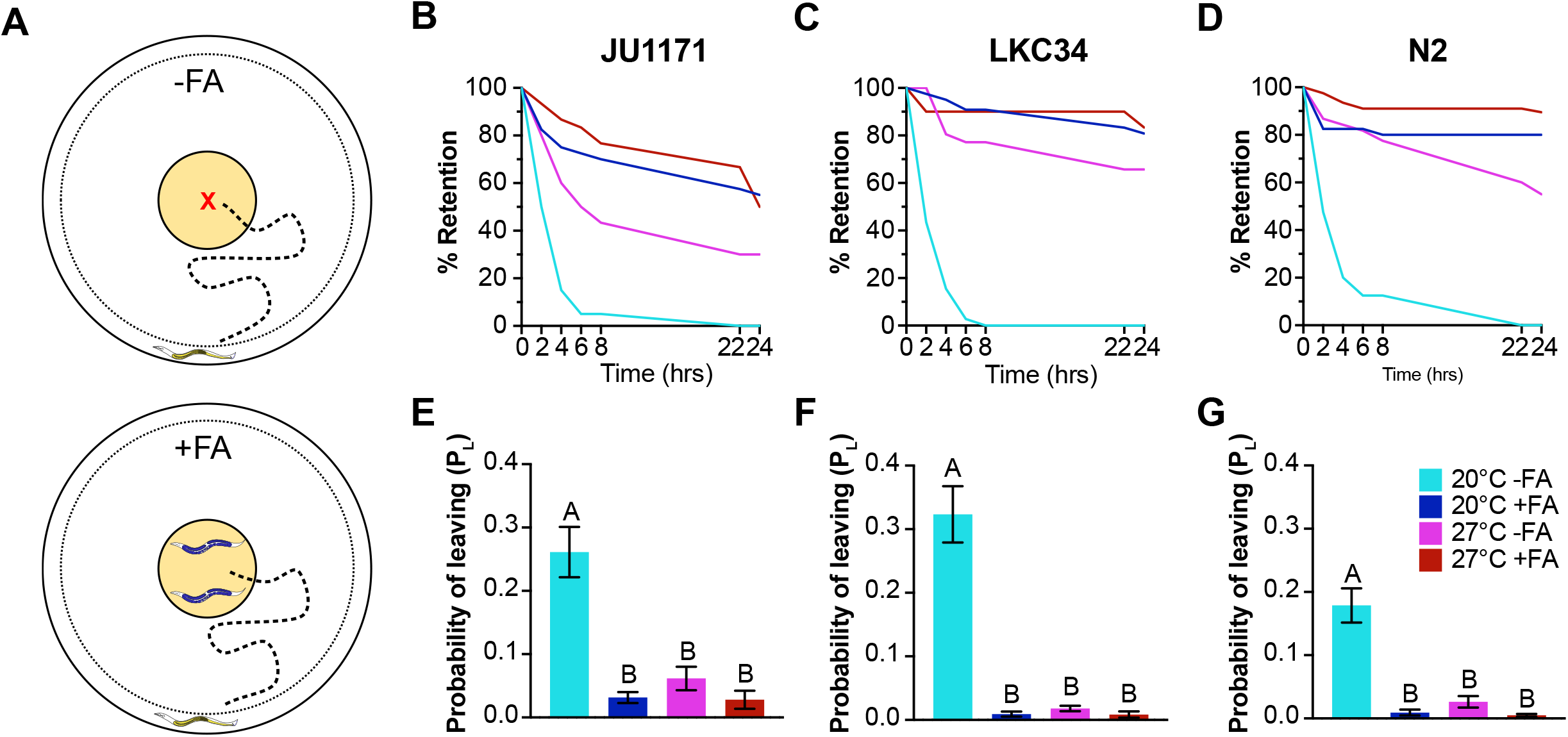
Males exposed to temperature stress rarely leave a food source whether hermaphrodites are present or not. (A) Schematic of the leaving assay. NGM plates are spotted with AMA1004 *E. coli* in LB. Control plates have no formaldehyde fixed hermaphrodites (FA hermaphrodites) (top). Test plates have two FA hermaphrodites placed on the food (bottom). Males were released on food (red x) and males are considered “leavers” if they wander within 2.5 mm of the edge of the plate (dotted circle) during the 24-hour experimental time frame. (B-D) Temperature stressed males rarely leave a food source regardless of the presence of FA hermaphrodites; however, unstressed males quickly leave a food source and engage in mate searching behavior in the absence of FA hermaphrodites. (E-G) Correspondingly, temperature stressed males have very low probabilities of leaving (P_L_) whether FA hermaphrodites are present or not. In contrast, unstressed males have high P_L_ values in the absence of FA hermaphrodites, and low P_L_ values in their presence. (Two-way ANOVA with Tukey’s correction, data with different letters are significantly different, data with matching letters are statistically identical. P value ≤ 0.05 Error bars represent ± 1 SEM).

### Exposure to 27°C may increase food drive in *C. elegans* males

Because temperature stressed males rarely leave a food source even in the absence of hermaphrodites (Fig. 2), we hypothesized that they have increased food drive in comparison to their unstressed counterparts. To test this possibility directly, we performed food search assays (Fig. 3A) (Ryan et al., 2014). We found that temperature stressed males from all three wild isolates are just as likely to reach a food source within the experimental time frame as their unstressed counterparts (Fig. 3 B-D). Because temperature stressed males have decreased motility compared to their unstressed counterparts (Fig. S1), but still reach food in a timely manner, we interpret these results to mean that exposure to 27°C increases food drive in *C. elegans* males.

**Fig. 3.**
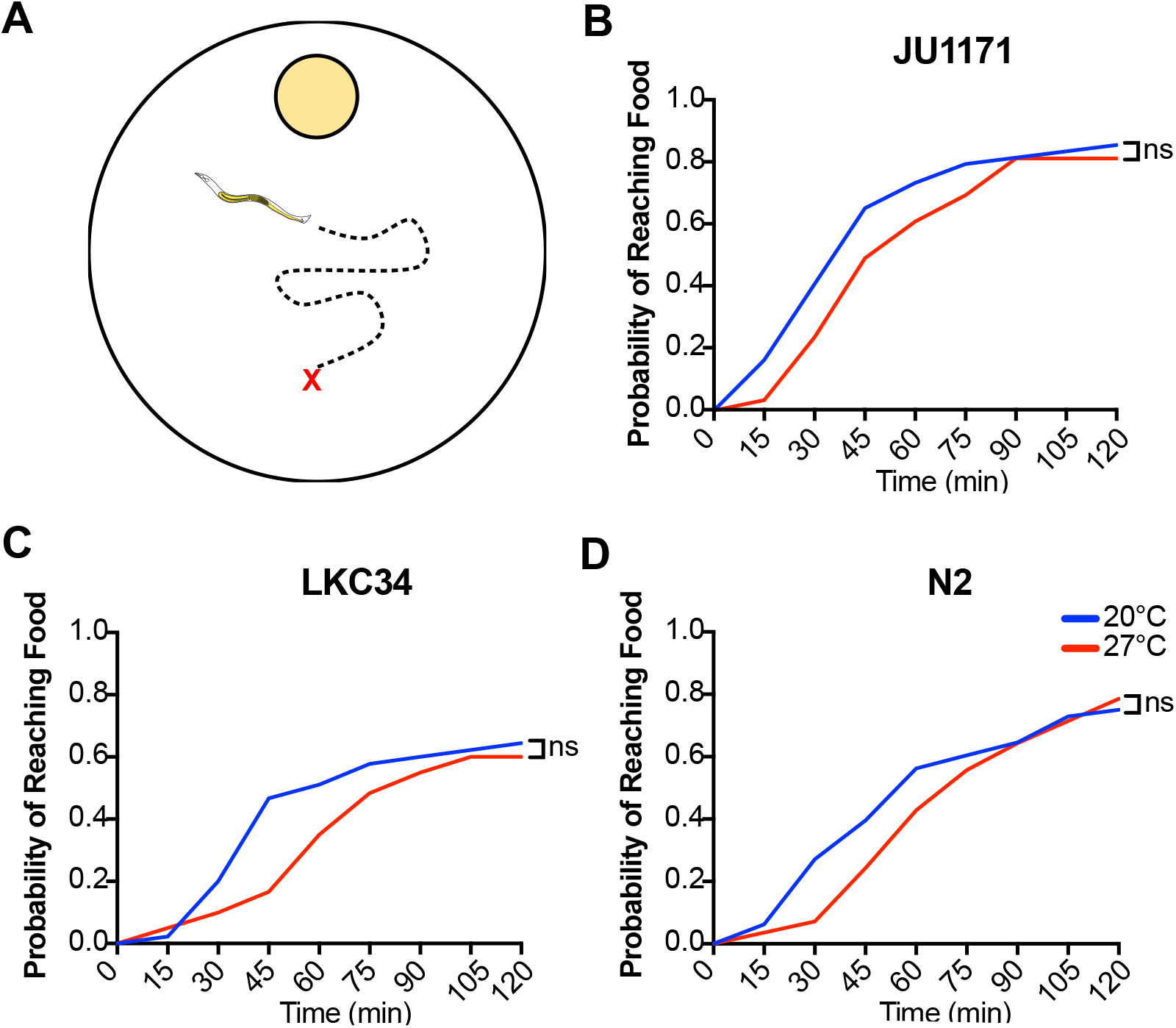
Temperature stressed *C. elegans* males are as likely to reach a food source during the experimental time frame as their unstressed counterparts. (A) Schematic of the food search assay. NGM plates are spotted with AMA1004 *E. coli* in LB (yellow). Males were released on the agar surface (red x) and observed every fifteen minutes for two hours to ascertain if they had reached the food spot. (B-D) Temperature stressed males from all three strains are equally likely to locate a food source as their unstressed counterparts (Log-rank [Mantel-Cox] test, ns indicates: P > 0.05).

### Temperature stressed male retention on food is not due to entrapment by physical properties of the food

Males exposed to 27°C rarely engage in mate searching behavior and leave a food source (Fig. 2). One possible explanation is that these males are unable to leave a food source because their locomotion defects (Fig. S1) result in their being trapped due to viscosity or other physical properties of the food. We tested this possibility by placing males on polystyrene microspheres, which mimic food viscosity without having any nutritional content or chemical signals. These experiments are identical to food search assays (Fig. 3A) except polystyrene microspheres are dispensed at the site where males are released (Fig. S2A). In high enough concentrations microspheres can be used to fully immobilize *C. elegans* and are often employed in microscopy in lieu of pharmacological paralytic agents (Manjarrez and Mailler, 2020). Additionally, they may be ingested by worms but provide no nutrition to the animal (Fueser et al., 2019, 2020, 2021). We used a concentration that does not fully immobilize worms but creates increased movement difficulty, similar to a bacterial lawn. We found that virtually all temperature stressed males leave the region with microspheres within the first fifteen minutes of the experiment and subsequently reach a food source similarly to males not exposed to microspheres (Fig. S2B-D). These results suggest that retention on a food source seen in temperature stressed males is not due to these males being trapped by the food source itself.

### Exposure to temperature stress compromises *C. elegans* male mate searching drive

Another possible explanation of diminished leaving behavior in temperature stressed males could be decreased mating drive (Barrios, 2014). To test this directly, we performed mate search assays (Fig. 4A) (Fagan et al., 2018) using two different types of hermaphrodites: Formaldehyde-fixed (FA) strain matched hermaphrodites that can be recognized by touch but do not produce any chemical signaling molecules, and live *unc-13(e51)* hermaphrodites in an N2 background that have a strong uncoordinated (Unc) phenotype and therefore do not move away from males, but still excrete chemical signaling molecules. We found that males of all three wild isolate strains were significantly less likely to locate FA or Unc hermaphrodites when exposed to temperature stress than their unstressed counterparts (Fig. 4B-D). However, we also found that males of all three wild isolate strains were significantly more likely to locate an Unc hermaphrodite than an FA hermaphrodite at both 20°C and 27°C (Fig. 4B-D). Collectively, these results suggest that temperature stress negatively impacts mate searching drive in *C. elegans* males, and that the chemical signals produced by hermaphrodites increase the likelihood that they will be found by searching males during the experimental time period at both 20°C and 27°C.

**Fig. 4.**
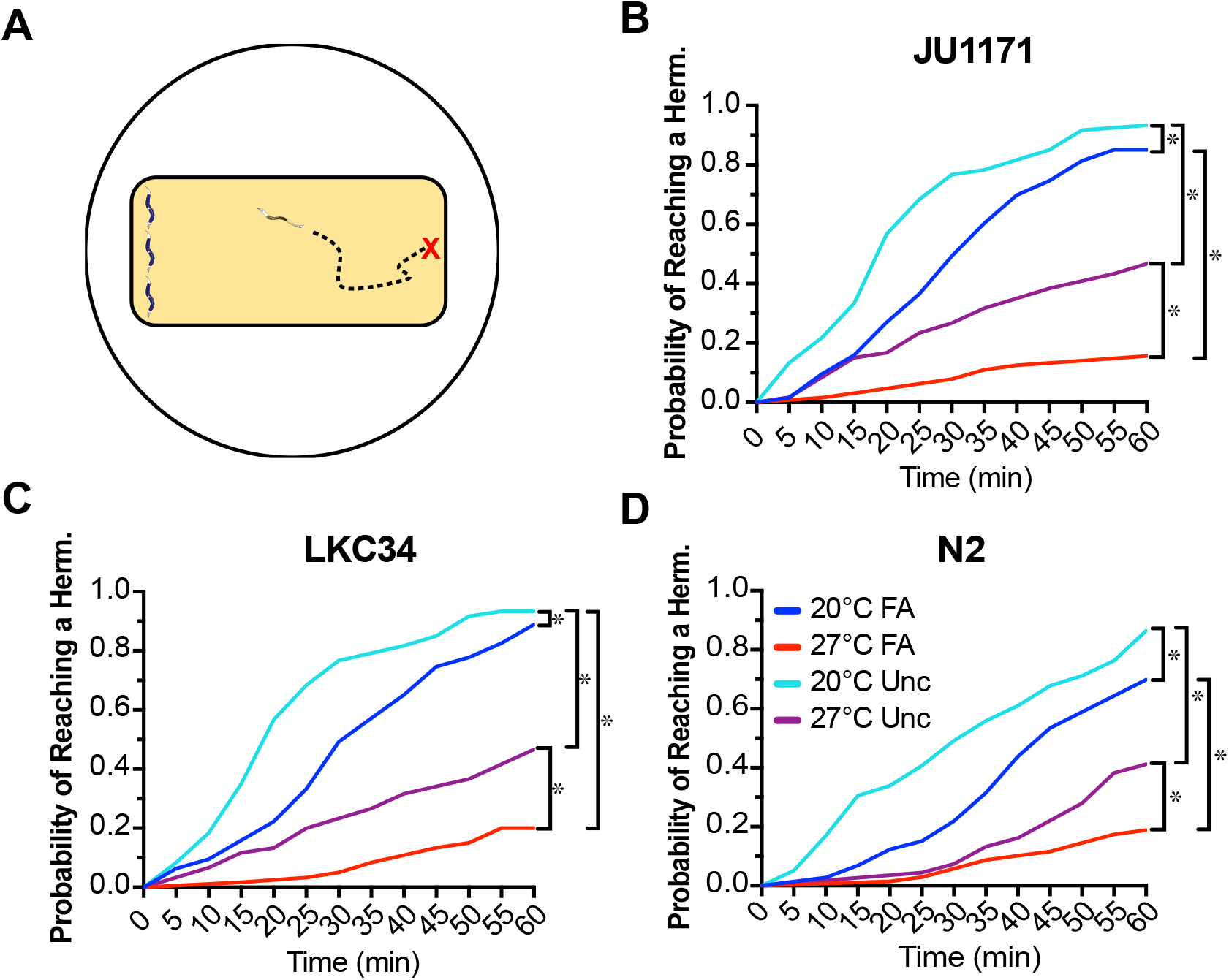
*C. elegans* males exposed to temperature stress are less likely to locate both formaldehyde-fixed (FA) and live Unc hermaphrodites. (A) Schematic of mate search assay. NGM plates were spread with AMA1004 *E. coli* in LB into a rectangular lawn (yellow). Formaldehyde fixed (FA) or live *unc-13(e51)* hermaphrodites were placed at one end of the lawn. Males were released on the opposite end of the lawn (red x) and observed every five minutes for one hour to ascertain if they had reached and responded to the hermaphrodites. (B-C) JU1171, LKC34, and N2 males raised at 27°C are significantly less likely to find either a FA or Unc hermaphrodite than their counterparts raised at 20°C. In addition, males from all three strains are more likely to locate a living Unc hermaphrodite than a FA hermaphrodite at either 20°C or 27°C (Logrank [Mantel-Cox] test, *P < 0.05).

### Temperature stressed males prioritize food over reproduction

The results of our previous experiments suggested that males may prioritize food over reproduction; opting to locate a food source and remain there whether hermaphrodites are present or not instead of engaging in mate-searching behavior. To test this possibility more directly, we designed a Food/Mate prioritization assay where males would be given the choice of a food spot without FA hermaphrodites and a food spot with FA hermaphrodites (Fig. 5A). We found that temperature stressed JU1171, LKC34 and N2 males randomly selected a food spot, and switched locations very infrequently whether on test plates (one food spot with FA hermaphrodites and one spot with no FA hermaphrodites) or control plates (two food spots with no FA hermaphrodites) (Fig. 5B-D). Unstressed males also randomly selected an initial food spot, however their subsequent behavior contrasts starkly with temperature stressed males. On test plates, males of all three strains would quickly accumulate on the spot with FA hermaphrodites and remain there, however, on control plates males would frequently switch between food spots (Fig. 6B-D). These results suggest that exposure to temperature stress reduces mating drive and increases food drive in *C. elegans* males.

**Fig. 5.**
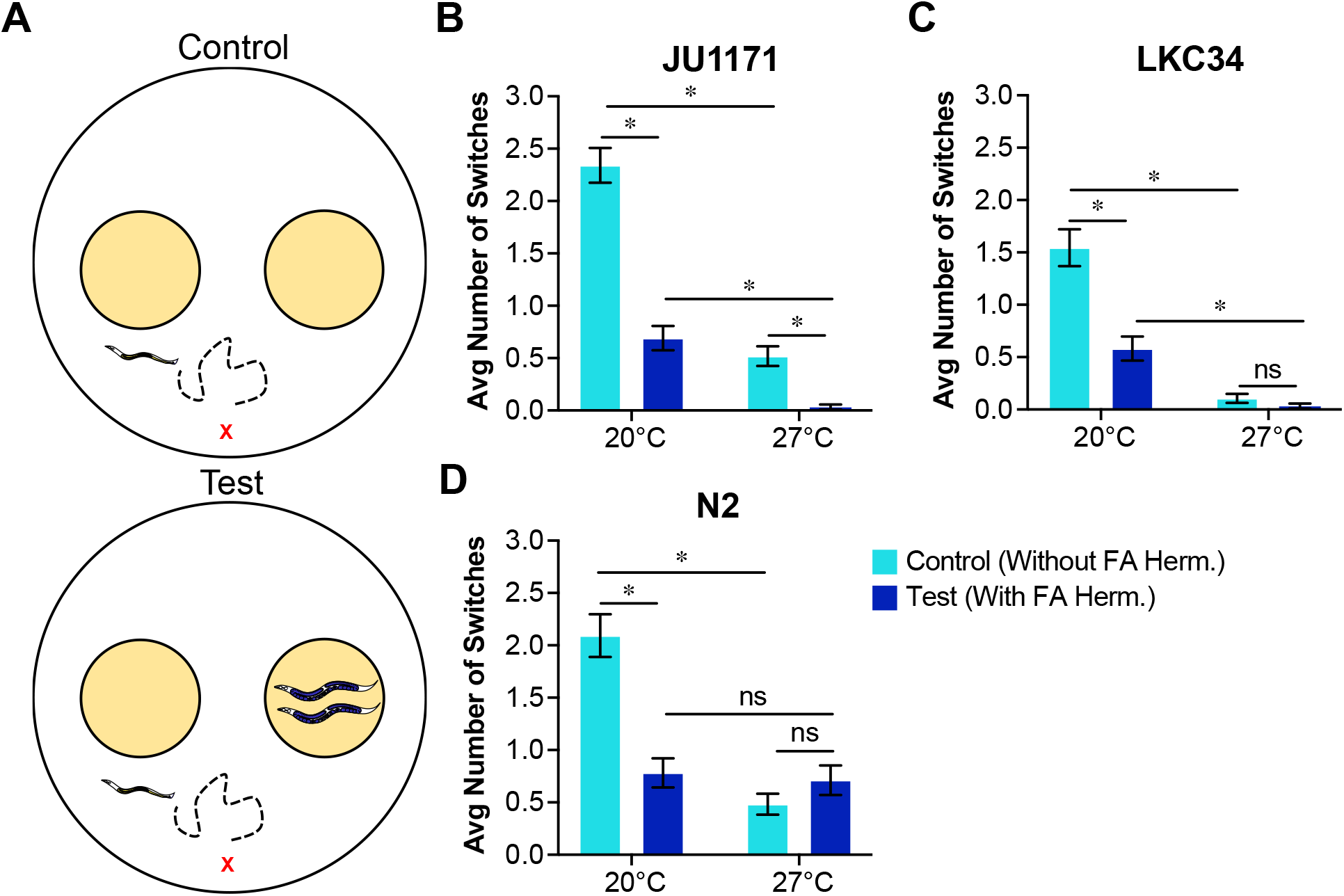
Temperature stressed *C. elegans* males are less likely to explore in search of a mate than their unstressed counterparts. (A) Schematic of the food/mate prioritization assay. Top: Control plates are set up with two AMA1004 *E. coli* in LB food spots but no FA hermaphrodites Bottom: Test plates are set up with two AMA1004 *E. coli* in LB food spots, one having two formaldehyde-fixed (FA) hermaphrodites. Males were released on the plate (red x) and their position was recorded during the 24-hour experimental time frame at the same time point as the leaving assay. (B) Temperature stressed JU1171 males are less likely to switch position than their unstressed counterparts whether placed on test plates or control plates. In addition, only temperature stressed JU1171 males are more likely to switch position on control plates that test plates (Two-way ANOVA with Tukey’s correction, *P ≤ 0.05). (C) Temperature stressed LKC34 males are less likely to switch position on either test plates or control plates than their unstressed counterparts (Two-way ANOVA with Tukey’s correction, *P ≤ 0.05). (D) Temperature stressed N2 males are less likely to switch positions on control plates than their unstressed counterparts but they are equally likely to change positions on test plates as their unstressed counterparts (Two-way ANOVA with Tukey’s correction, ns > 0.05). Error bars represent ± 1 SEM.

**Fig. 6.**
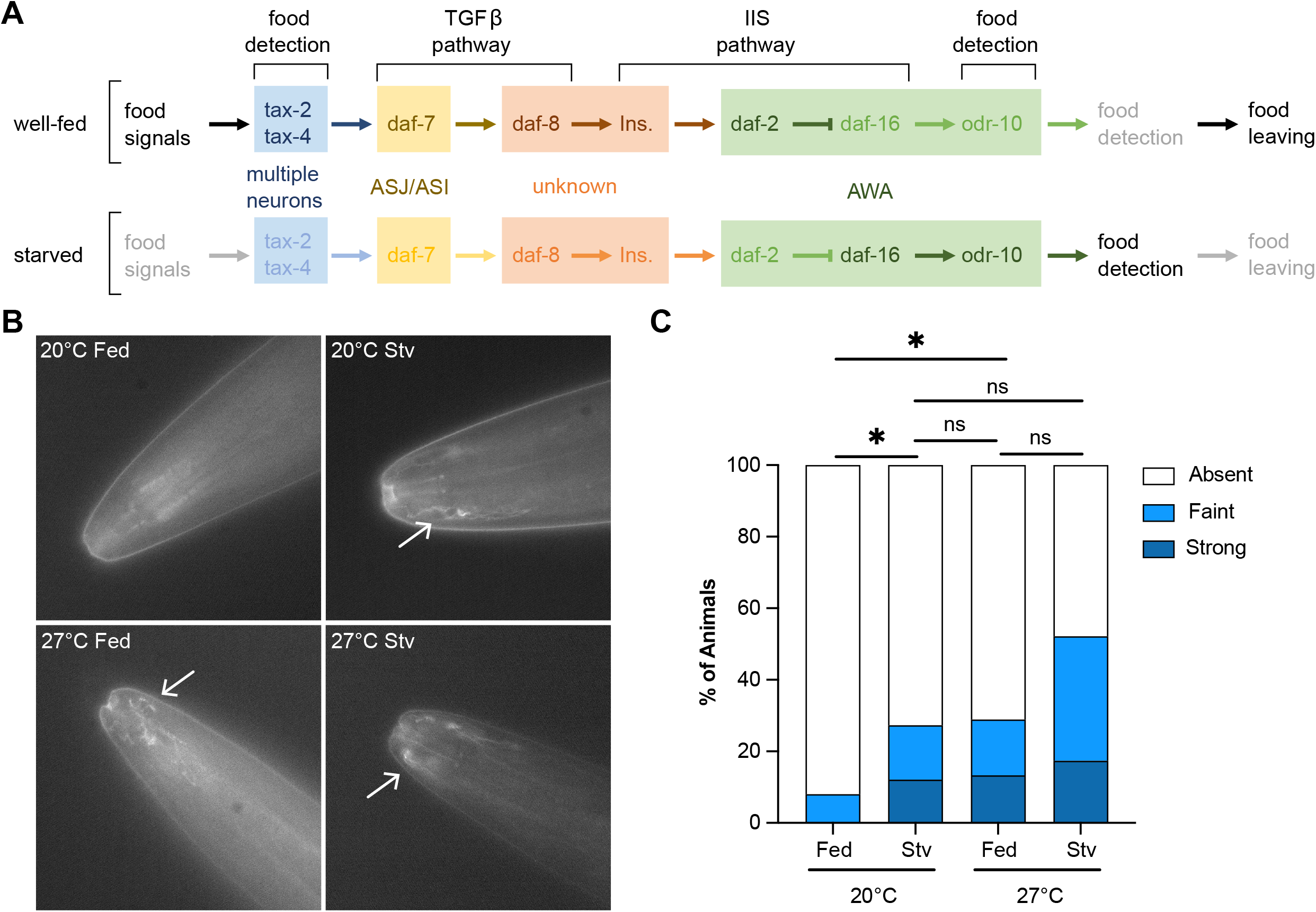
Both well-fed and starved males raised at 27°C show increased levels of ODR-10 expression. (A) Schematic showing the regulation of *odr-10* by nutritional state in *C. elegans* males (Wexler et al., 2020). (B) Males were raised at either 20°C or 27°C and starved or well-fed from young adulthood overnight before ODR-10::GFP fluorescence was scored. Representative micrographs showing ODR::GFP: (top left) a well-fed male at 20°C, (top right) a starved (stv) male at 20°C, (bottom left) a well-fed male at 27°C, and (bottom right) a starved (stv) male at 27°C. (C) The proportion of temperature stressed males, both well-fed and starved, with ODR-10::GFP fluorescence was higher than well-fed males at 20°C but was the same as starved males at 20°C (Chi-square test on 2X3 contingency table comparing temperature/nutritional status by fluorescent intensity, *P ≤ 0.05).

### Temperature stressed males have increased ODR-10::GFP expression

We observed that males exposed to 27°C reach a food source in a timely manner (Fig. 3B-D), and very rarely leave a food source even when hermaphrodites are absent (Fig. 2B-D) even when given the option of a food source with hermaphrodites (Fig. 5B-D). This led us to believe that temperature stressed males may have an increased food drive due to their changes in their ability to detect and respond to signals from food. *C. elegans* males ascertain their nutritional status via external chemical cues from a food source (Wexler et al., 2020). In well-fed unstressed males, food signals increase the expression of *daf-7/TGFβ* which suppresses expression of the food associated chemoreceptor *odr-10* (Fig. 6C), promoting leaving behavior (Wexler et al., 2020). In starved unstressed males, *odr-10* expression is high (Fig. 6A), prompting a return to feeding behavior (Wexler et al., 2020). We hypothesized that ODR-10 levels are higher in temperature stressed males, promoting feeding behavior and retention on a food source. In order to test this possibility we examined males carrying the ODR-10::GFP transgene *kyIs53 [Podr-10::odr-10::GFP]*. We found that well-fed temperature stressed males have higher levels of ODR-10::GFP than their well-fed unstressed counterparts (Fig. 6B). In fact well-fed temperature stressed males have levels of ODR-10::GFP equivalent to both well-fed and starved unstressed males (Fig 6B). These data suggest that temperature stress increases the amount of ODR-10 in male worms which may subsequently promote male feeding behavior and retention on a food source.

## Discussion

In the research presented here, we endeavored to identify the reasons why *C. elegans* males exposed to moderate temperature stress (27°C) have little to no interest in mating. A male uninterested in mating with a hermaphrodite stands virtually no chance at reproducing successfully, regardless of the quality of his sexual anatomy or germline-associated processes like sperm production and quality. We found that temperature stressed males lack interest in hermaphrodites due to an upset in the delicate balance of their priorities: mating and feeding. Temperature stressed *C. elegans* males prioritize feeding over mating to such extent that they are virtually sterile at 27°C.

### Temperature stressed *C. elegans* male food prioritization is due to perceived hunger

*C. elegans* males have two priorities: (1) feeding, and (2) mating (Barrios, 2014). The ideal scenario for a male is a source of plentiful food with numerous potential mates. We were surprised to discover that temperature stressed males rarely leave a food source in the absence of hermaphrodites, and when presented with an ideal scenario (food and mates overlapping spatially and temporally) they very rarely approach and copulate with hermaphrodites. Equally surprising, temperature stressed males locate a food source as quickly as their unstressed counterparts despite their considerable motility defects. Thus, temperature stressed males appear to have a reduced mating drive and an increased food drive. *C. elegans* male exploratory behavior is regulated by both internal states and sensory cues (Barrios, 2014), any or all of which could be temperature sensitive.

We looked at one such state shown to regulate exploratory behavior: Nutritional status. Starved males temporarily prioritize feeding until they are well-fed, after which normal exploratory behavior resumes in in the absence of hermaphrodites at 20°C (Lipton et al., 2004). *C. elegans* males are satiated by the perception of food, rather than internal cures elicited by ingestion of bacteria (Wexler et al., 2020). In food-deprived male worms, a lack of chemical signals from food leads to reduced *daf-7 (C. elegans* TGF-β) expression in the ASJ and ASI sensory neurons and a subsequent decline in insulin signaling via *daf-2 (C. elegans* insulin receptor) in the AWA sensory neurons (Wexler et al., 2020). Resultant increases in *daf-16 (C. elegans* FOXO transcription factor) expression AWA, activates the food signal chemoreceptor *odr-10* in AWA (Wexler et al., 2020). Increased levels of ODR-10 are associated with an increase in feeding behavior in starved males; and upon feeding ODR-10 levels return to baseline (Ryan et al., 2014; Wexler et al., 2020). Because well-fed males at 27°C show levels of ODR-10 expression equivalent to starved males at 20°C, we propose that temperature stressed males perceive themselves to be starving due to food signals not being received nor interpreted when food is present, promoting feeding behavior over mate seeking behavior. It has been shown that chemoreceptor expression in *C. elegans* is regulated by various internal and external cues (Peckol et al., 2001; Lanjuin and Sengupta, 2002; Ryan et al., 2014; Wexler et al., 2020), but to our knowledge this is the first example of chemoreceptor expression regulated by elevated temperature in *C. elegans*.

### The nervous system of *C. elegans* males raised at 27°C may exhibit broad changes in function

Contrary to our initial prediction, *C. elegans* males exposed to temperature stress remain in proximity to hermaphrodite chemical signals for longer than their unstressed counterparts, even though there are no hermaphrodites physically present in the area. This change in behavior could be considered maladaptive. *C. elegans* male attraction to hermaphrodite excreted chemical signals could improve their chances of securing a mate, however lingering in an area with these signals despite a lack of physical contact with hermaphrodites would squander both time and energy. This inability to make the decision to leave a region conditioned by hermaphrodites with could be due to changes in structure or function of the sensory neurons that receive these chemical signals, which include both sex specific (CEM) and shared (AWA, AWC, ADF, ASK) neurons (Chute and Srinivasan, 2014; Muirhead and Srinivasan, 2020). Interestingly, pheromone, food, and temperature sensing connect in a shared circuit. The AWA sensory neurons mediate male attraction to both ascaroside pheromones (White et al., 2007) and volatile sex pheromones (Wan et al., 2019). In addition AWA are the only known synaptic input to the AFD thermosensory neurons (Mori and Ohshima, 1995; Kimura et al., 2004; Clark et al., 2006; Ramot et al., 2008), which are essential for *C. elegans* thermotaxis (Goodman and Sengupta, 2018, 2019). AWA are also the site of *odr-10* expression and function, which mediates food attraction and where we see changes in *odr-10* regulation (Wexler et al., 2020). AWC are another set of sensory neurons that play a role in male attraction to ascarosides (White et al., 2007), as well as their role in thermosensation (Biron et al., 2008; Kuhara et al., 2008). Finally, ASI chemosensory neurons are an important component of the thermosensory circuit (Beverly et al., 2011) and the site of *daf-7* expression, which regulates *odr-10* in the AWA neurons and thus food attraction (Wexler et al., 2020). It seems altogether possible that temperature stress has a broad excitatory effect on these thermosensitive neurons. For example, increased activity in the AWA neurons due to temperature stress could affect male behavioral response to food, and reproductive signals. Due to increased or aberrant activity in these neurons, temperature stressed males may be unable to decide leave a food source, or a region conditioned by hermaphrodites despite a distinct lack of potential mates.

We have also shown previously that males exposed to temperature stress at some point in their lives will complete mating without actually transferring sperm (Nett et al., 2019) or ejaculate on the hermaphrodite’s body wall instead of in the reproductive tract (N.B.S. unpublished observations). Difficulties with sperm transfer are a hallmark of changes in excitability in the reproductive circuit of senescent *C. elegans* males (Guo et al., 2012; Guo and García, 2014). *C. elegans* males use copulatory spicules and associated muscles in their tail to breach the hermaphrodite vulva and transfer sperm (Barr et al., 2018). As males age, the sex neuromusculature becomes hyperexcitable, impeding sperm transfer and ultimately leading to significantly reduced fertility (Guo et al., 2012; Guo and García, 2014). In addition to advanced age, elevated temperature has been shown to broadly increase neuronal hyperexcitability in *Drosophila* (Peng et al., 2007), mice (Graham et al., 2008), and humans (Mizunuma et al., 2009). If this is the case in temperature stressed *C. elegans* males, defects in mating behaviors may be due in part to widespread changes in neuronal excitability at 27°C. In order to address these possibilities directly, the appropriate sensory neurons’ structure and function should be addressed using in vivo neuronal calcium imaging (Chung et al., 2013; Nguyen et al., 2016; Venkatachalam et al., 2016).

### Differences between wild isolate strains may be relevant in their natural environments

A majority of *C. elegans* research is confined to the laboratory domesticated canonical wild type strain N2, with comparatively fewer studies exploiting the growing number of wild isolate strains which more fully capture the genetic diversity of the species (Andersen et al., 2012; Sterken et al., 2015; Cook et al., 2017). Fewer studies still are performed with wild isolate males. We’ve shown elsewhere that there are appreciable differences between wild isolate male fertility under temperature stress (Petrella, 2014; Nett et al., 2019; Sepulveda and Petrella, 2021), and in this study we’ve dissected fertility into several behavioral factors which also differ between isolates. The question remains however, how relevant are these differences in the wild? *C. elegans* appears to reproduce preferentially in rotting fruit and herbaceous plant matter at the surface where bacteria are plentiful, but where temperatures can reach those that we use in our experiments (Kiontke et al., 2011; Félix and Duveau, 2012; Petersen et al., 2015; Richaud et al., 2018). Males are present in the wild at low levels and outcrossing appears to be infrequent in both ideal environs like rotting fruit (Richaud et al., 2018) and less desirable locations like compost heaps (Barrière and Félix, 2005, 2007; Haber et al., 2005; Sivasundar and Hey, 2005), possibly due to outbreeding depression (Dolgin et al., 2007). However, a single migratory male from a strain that fares comparatively well under temperature stress, like JU1171, could fare very well in a population of hermaphrodites where he is faster, has more physical endurance, possesses a less hyperexcitable mating circuit, and transfers sperm more reliably than a different genotype of male if there exist no genetic incompatibility between the strains (Seidel et al., 2008, 2011). While evidence of outcrossing suggests it is infrequent in many natural populations of *C. elegans*, several studies have demonstrated that it is adaptive in novel and stressful environments (Morran et al., 2009a, 2009b; Teotonio et al., 2012; Carvalho et al., 2014; Chelo et al., 2014). Climate change associated changes in ambient temperature may prove to be on such novel and stressful environment that favors *C. elegans* males, especially those with superior reproductive features under temperature stress.

## Fig. Legends

**Fig. S1.**
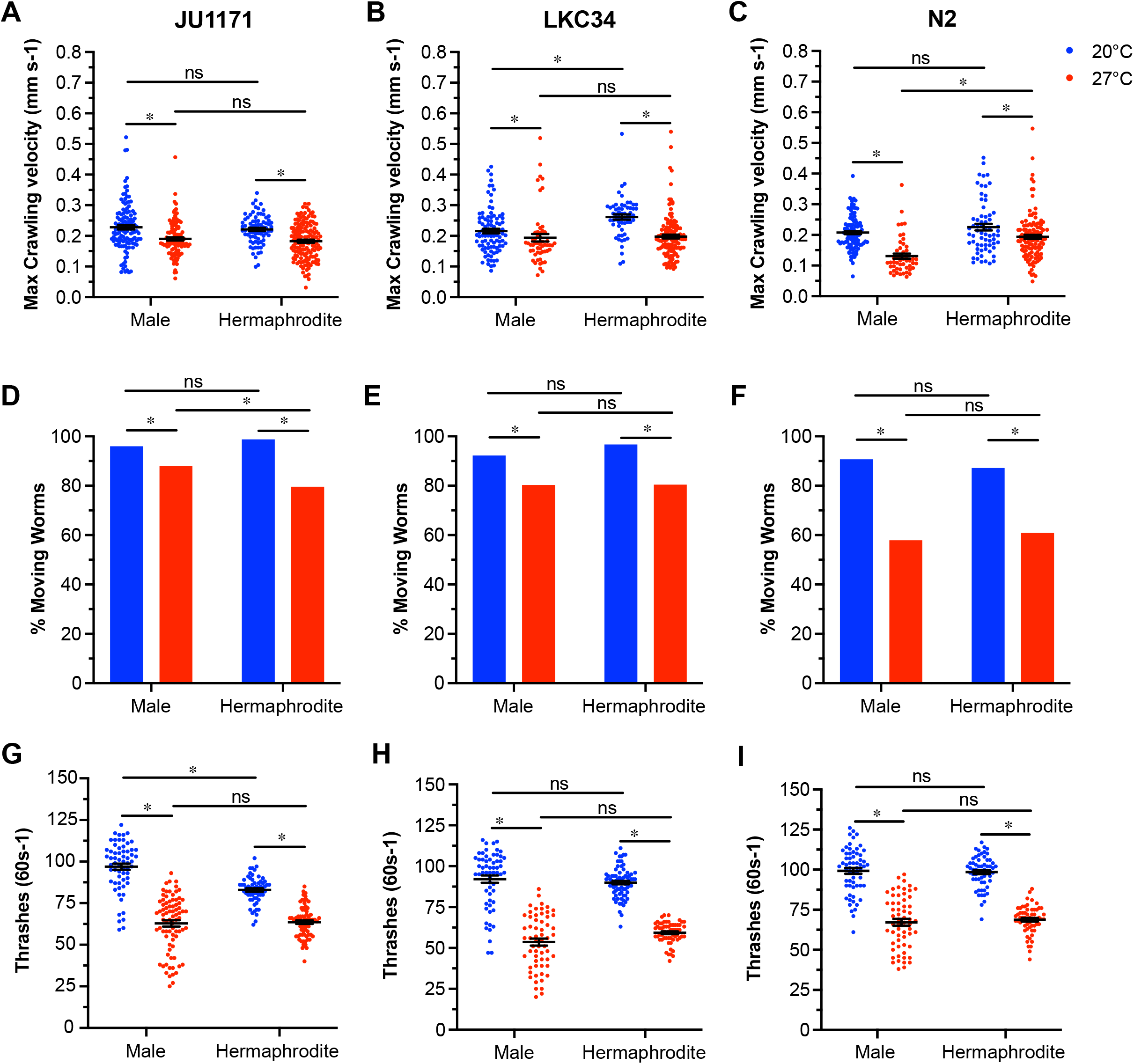
Upon exposure to temperature stress, *C. elegans* males and hermaphrodites have lower maximum crawling velocities and reduced physical endurance. (A-C) JU1171, LKC34, and N2 males and hermaphrodites were acclimated on an NGM plate with a thin layer of food and videos were recorded. The videos were analyzed to determine their maximum crawling velocities. Males and hermaphrodites raised at 27°C have lower maximum crawling velocities than their counterparts raised at 20°C. LKC34 hermaphrodites have higher maximum crawling velocities than males at 20°C. N2 hermaphrodites have higher maximum crawling velocities than males at 27°C (Two-way ANOVA with Tukey’s correction, *P ≤ 0.05). (D-F) The proportion of JU1171, LKC34, and N2 males and hermaphrodites that moved on an NGM plate with a thin layer of food was also scored. JU1171, LKC34, and N2 males and hermaphrodites raised at 27°C are less likely to be motile than their counterparts raised at 20°C. JU1171 males move more at 27°C than hermaphrodites (Fisher’s Exact Test, *P < 0.05). (G-H) JU1171, LKC34, and N2 males and hermaphrodites were placed in liquid buffer and the number of full body thrashes was scored over one minute to assess physical endurance. JU1171, LKC34, and N2 males and hermaphrodites raised at 27°C have less physical endurance (perform fewer full body thrashes) than their counterparts raised at 20°C. JU1171 males are more vigorous than hermaphrodites at 27°C (Two-way ANOVA with Tukey’s correction, *P < 0.05).

**Fig. S2.**
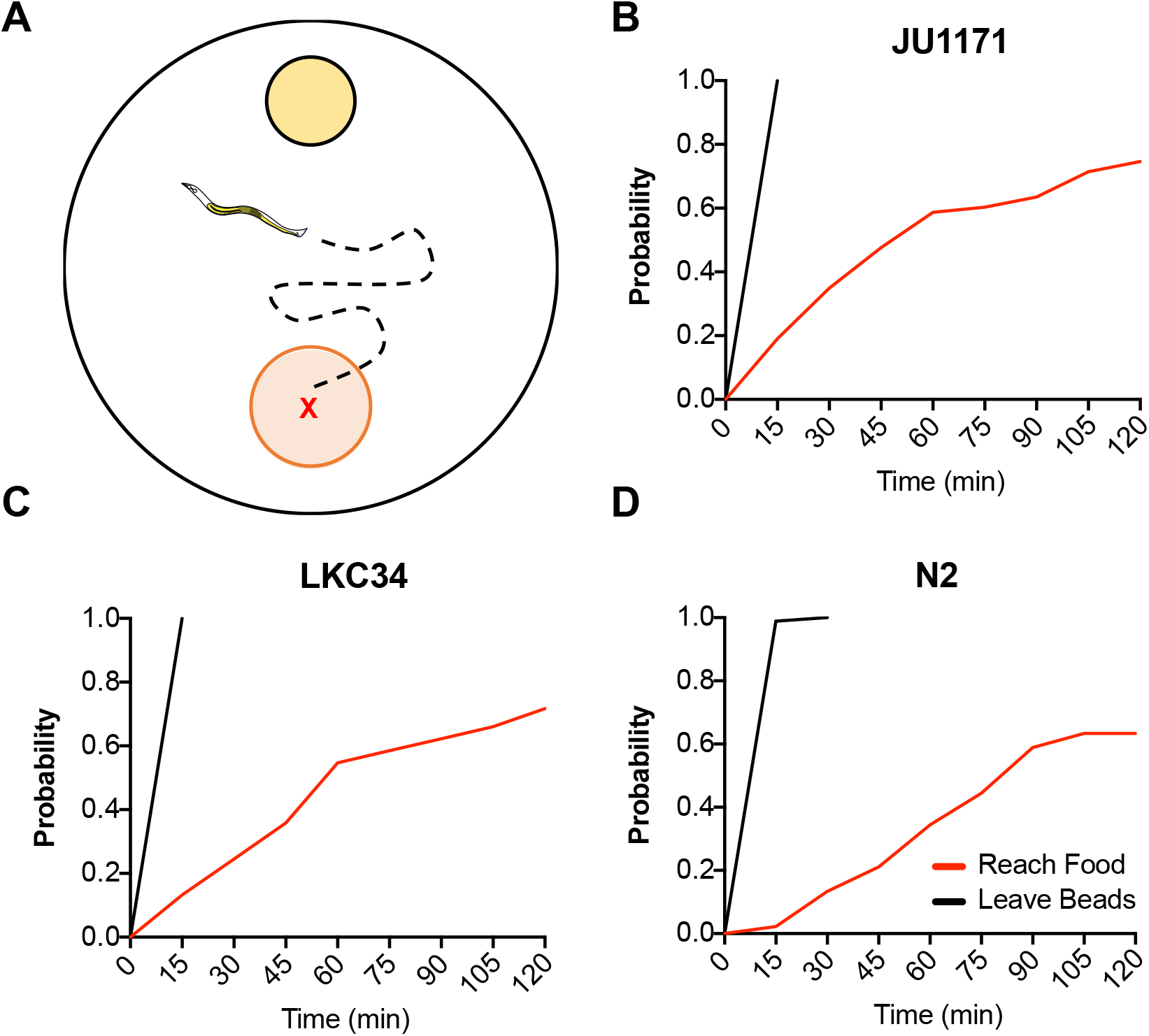
Temperature stressed males are not trapped by polystyrene microspheres. (A) Schematic of the Polystyrene microsphere immobilization assay. NGM plates were spotted with AMA1004 *E. coli* in LB (yellow). Microspheres were dispensed on the agar surface (red). Males were released in the center of the microspheres (red x) and observed every fifteen minutes for two hours to ascertain if they had escaped the microspheres and reached the food spot. (B-D) JU1171, LKC34, and N2 males exposed to temperature stress quickly leave the region spotted with microspheres and locate a food source similarly to counterparts not placed on microspheres (Log-rank [Mantel-Cox] test, ns indicates: P > 0.05).

---

**Table 1:** Strains used in study.

| **Strain** | **Genotype** | **Genetic Background** | **Available from** |
| --- | --- | --- | --- |
| DR476 | *daf-22(m130)* II | N2 | CGC |
| EM930 | *daf-9(m540)* X;<br>*bxls14*, *him-5(e1490)*<br>V | N2 | Emmons lab |
| JU1171 | Wild isolate | JU1171 | CGC |
| LKC34 | Wild isolate | LKC34 | CGC |
| MT7929 | *unc-13(e51)* I | N2 | CGC |
| N2 | Lab wild type | N2 | CGC |
| UR460 | *him-5(e1490)* V;<br>*kyls53* [*Podr-10::odr-10::GFP*] X | N2 | Portman lab |

